# Withdrawal latency in a human hot plate analog tracks inter-individual differences in pain sensitivity

**DOI:** 10.64898/2026.10.16.759590

**Authors:** Lars Keuter, Ole Goltermann, Jonathan Jeßberger, Leonard Asan, Christian Büchel

## Abstract

Pain research relies mainly on self-report in humans and nocifensive behavior in animals. This asymmetry limits cross-species translation and neglects movement as integral to human pain responses. We therefore developed a human analog of the rodent hot plate test and tested whether temporal and kinematic features of nocifensive withdrawal can capture inter-individual differences in pain sensitivity. Ninety-one healthy participants rested their forearm on a thermode that rapidly heated to and held a fixed noxious temperature (46 °C or 48 °C) and were instructed to withdraw once the percept reached their pain threshold or tolerance. From wrist accelerometry, we extracted withdrawal latency and peak acceleration, an index of withdrawal vigor, and related both to pain intensity ratings from a separate, temperature-matched task and to calibrated pain sensitivity. Participants withdrew earlier and with higher peak acceleration at 48 °C than at 46 °C. Crucially, withdrawal latency tracked inter-individual differences in pain sensitivity: Participants who withdrew earlier reported higher pain in the separate task (48 °C: *r* = −.47 and −.50) and showed lower pain thresholds and higher suprathreshold sensitivity during calibration. In contrast, withdrawal vigor showed no such relationship, despite high reliability. Withdrawal latency can thus serve as a behavioral readout of inter-individual differences in pain sensitivity, whereas vigor might reflect intra-individual motor scaling with stimulus intensity. By separating the decision to withdraw from its motor execution, this paradigm offers a behavioral complement to traditional self-report and provides a translational link between human and preclinical pain assays.

## Introduction

Pain research in humans and animal models relies on fundamentally different measures. While human pain is commonly assessed through self-report, pain in non-human animals is inferred primarily through nocifensive behaviors such as withdrawal [8]. This methodological asymmetry hinders the translation of findings between species. It also leaves a critical channel underexplored: Although movement is an integral part of the human response to noxious stimulation [23], it is rarely used to quantify pain or inter-individual differences in pain sensitivity.

In preclinical assays, nocifensive motor behavior is the primary readout of nociception. In the hot plate test, a freely moving animal is placed on a surface held at a noxious temperature and the latency to a nocifensive behavior is recorded [8,31]. Efforts to translate these behavioral readouts for human use have been sparse but promising. Adapting the rodent tail-flick test, Lee and Stitzer [17] showed that human finger withdrawal latency to radiant heat remains stable across test sessions. This latency shortens as stimulus intensity increases and varies with the instruction given to participants [10]. More recently, Kamper-Fuhrmann and colleagues [14] extended the approach to contact heat, demonstrating that when participants self-terminated a ramped thermal stimulus by withdrawing their hand, the resulting pain threshold estimates were comparable to standard psychophysical methods.

These studies suggest that human withdrawal latency provides a reliable, intra-individual marker of nociceptive processing. However, it remains untested whether such behavior can capture inter-individual differences in pain sensitivity. Furthermore, classical assays typically reduce behavioral analysis to a single temporal dimension, focusing on *when* the movement occurs rather than *how* it is physically executed. The vigor with which a person performs a movement – indexed by metrics such as limb speed or acceleration – exhibits stable trait-like inter-individual differences [27]. Human pain sensitivity similarly varies robustly across individuals [7,15]. Because nocifensive withdrawal is a direct motor output of nociceptive processing, these inter-individual differences could manifest in both when and how the movement is performed. As such, both the timing (latency) and vigor (acceleration) of withdrawal may complement traditional self-report.

To investigate this, we developed a human analog of the hot plate test designed to capture withdrawal kinematics. Participants withdrew their forearm from a thermode maintained at a fixed noxious temperature, with steady baseline contact ensured via a load cell. Using a fixed temperature rather than a ramp ensures that all participants encounter the same target temperature, so that escape kinematics are not confounded with the temperature at withdrawal. Concurrently, a wrist-mounted accelerometer quantified the temporal and kinematic features of the withdrawal response. First, to test whether these metrics scale intra-individually with stimulus intensity, we compared withdrawal features between 46 °C and 48 °C. Furthermore, to test whether withdrawal features index inter-individual differences in pain sensitivity, we correlated both withdrawal latency and peak acceleration with thermal pain intensity ratings obtained from the same individuals (*N* = 91) in a separate, temperature-matched task within the same session. In addition, we related both features to the heat pain threshold and suprathreshold pain sensitivity obtained in a pre-task calibration.

## Materials and Methods

### Participants

The study was approved by the Ethics Committee of the Hamburg Medical Association (code 2024-101416-BO-ff) and conformed to the Declaration of Helsinki. All participants provided written informed consent. A total of 110 healthy participants were recruited through online advertisement (www.stellenwerk.de) and through word of mouth. The present experiment is part of a larger functional magnetic resonance imaging (fMRI) study for which the sample size was planned. Participants were screened for MRI- and study-specific exclusion criteria. Participants were excluded if they were under 18 or over 45 years old, had uncorrectable vision issues, or had MRI contraindications such as claustrophobia or a pacemaker. Further criteria were pregnancy, breastfeeding, self-reported chronic pain, or diagnosed neurological, psychiatric, dermatological, or inner-ear conditions. Participants were also excluded if they were enrolled in a pharmacological study, took regular medications, or had taken analgesics in the 24 hours before the experiment. Of the 110 participants recruited for the larger study, nine were excluded from the final sample because of thermode malfunctions (n = 7), excessive head motion during MRI acquisition (n = 1), or discontinuation of the session due to claustrophobia (n = 1). Of the remaining 101 participants, five were tested before the thermal withdrawal task had been finalized (without the load cell) and were excluded from the present analyses. A further five were excluded because of technical failures during the thermal withdrawal task. The final sample thus comprised 91 participants (47 female, 44 male; mean age = 27.7 years, SD = 6.2, range 20–44).

### Experimental procedure

The study comprised a single session, lasting approximately 3.5-4 hours, of which the thermal withdrawal task took approximately 30 minutes. Upon arrival, each participant provided written informed consent and was introduced to the experimental design and session structure. The session began with a calibration of each participant’s heat pain sensitivity on the left forearm. Participants then completed the thermal withdrawal task, followed by a thermal rating task inside the scanner, and finally T1-weighted and diffusion-weighted structural imaging. After exiting the scanner, participants completed a questionnaire battery (data not reported here) and were debriefed.

#### Apparatus

The thermal withdrawal task used an MRI-compatible contact heat-evoked potential stimulator thermode equipped with 30 × 30 mm² Peltier elements (CHEPS; Medoc, Israel), controlled from MATLAB 2023b through an Arduino-based interface. The thermode was mounted on a 3D-printed cast housing a load cell (Fig. 1A). The load cell signal was amplified and digitally converted by an HX711 at a sampling rate of 10 Hz and transmitted to a PC via an Arduino Nano. In addition, a tri-axial accelerometer (TSD109C2-MRI, Biopac Systems) was attached to the participants’ left wrist to record acceleration at 2,000 Hz. The left forearm rested on a cushion, so that the arm and thermode were level, with the accelerometer wristband positioned flush against the edge of the thermode to ensure consistent sensor placement across participants.

**Fig. 1.**
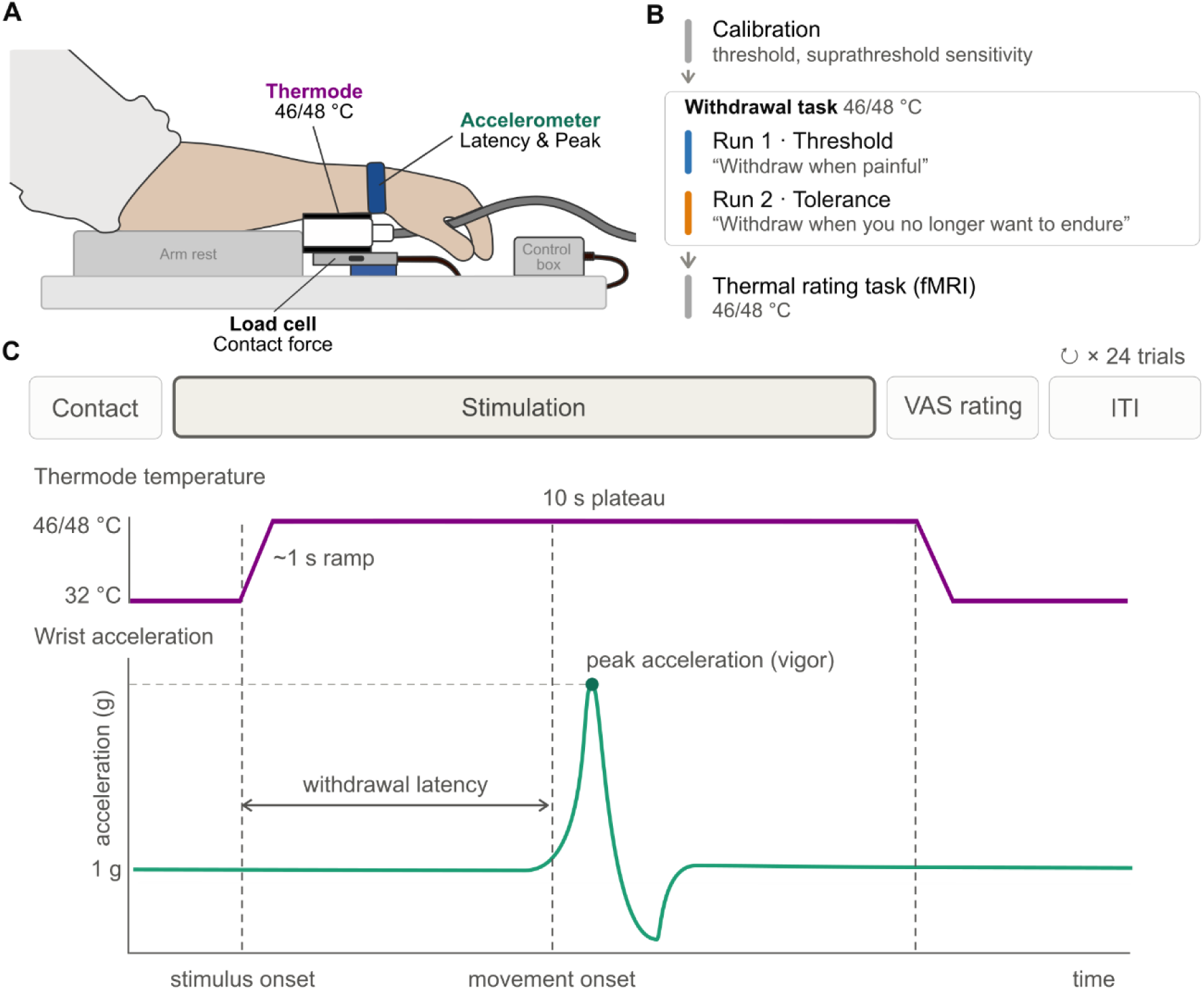
Experimental apparatus, session timeline, trial structure, and temporal profile of thermal stimulation and withdrawal response. **(A)** Schematic of the experimental apparatus. Participants placed their arm on an armrest with a thermode applied to the left forearm (with the accelerometer band aligned flush against the thermode, ensuring consistent placement). Contact force was monitored via an underlying load cell, and a wrist-worn accelerometer recorded withdrawal latency and peak acceleration (withdrawal vigor). **(B)** Session timeline. Heat pain threshold and suprathreshold sensitivity were calibrated on the left forearm before the withdrawal task. The withdrawal task comprised a pain threshold run (“withdraw when painful”) followed by a pain tolerance run (“withdraw when you no longer want to endure”), each with 46 °C and 48 °C stimuli. Pain ratings to the same temperatures were then obtained in a separate thermal rating task in the MRI scanner (left forearm and lower leg). **(C)** Trial structure and single-trial temporal profile. The top row illustrates the trial sequence, repeated for 24 trials (12 per run). Following confirmed steady contact, a red fixation cross cued stimulus onset. Participants withdrew their forearm based on the instructed criterion. After the thermode returned to the baseline temperature (32 °C), participants provided a pain intensity rating (0-100 VAS; 50 = minimally painful), followed by an 8-10 s inter-trial interval. The bottom row illustrates a representative accelerometer recording. The thermode temperature ramped from baseline to the target temperature, where it was held for a 10 s plateau. From the continuous acceleration recording, two features of the withdrawal were extracted: withdrawal latency (the time from stimulus onset to movement onset) and peak acceleration (the maximum acceleration of the withdrawal).

#### Thermal stimulation

The baseline temperature was set to 32 °C. On each trial the thermode ramped to a fixed target of 46 °C or 48 °C at 13 °C/s (ramp durations 1.08 s and 1.23 s), held the plateau for 10 s, and returned to baseline. Each run contained six trials per temperature, ordered pseudorandomly with no more than three consecutive trials at the same temperature.

#### Load cell calibration

At the beginning of each run, the load cell was tared with the arm off the thermode. Participants then placed the forearm on the thermode, and the resting contact force was sampled for 5 s. The mean ± 5 SD of that contact sample defined a participant- and run-specific acceptance window used for the contact check on all subsequent trials. The experimenter checked the resulting values and could accept them or repeat the calibration. Finally, the load cell was tared again.

#### Thermal withdrawal task

Participants placed their left forearm on the thermode so that the accelerometer Velcro strap and the thermode were aligned, ensuring comparable thermode positioning across participants. They then completed two runs of 12 trials. In the first run, participants withdrew the arm when the stimulus became painful (threshold instruction); in the second, when they no longer wanted to endure the pain (tolerance instruction; Fig. 1B). Threshold always preceded tolerance, so that threshold judgments could not be influenced by prior exposure to longer, tolerance-level stimulation (e.g., through sensitization or a shift in the internal criterion).

The trial structure is shown in Fig. 1C. Each trial began with the instruction to place the arm on the thermode. Contact was accepted once load cell readings fell inside the acceptance window continuously for 0.5 s. The experimenter had the option to re-tare or to confirm contact manually. After a 1 s delay, heat onset was signaled by a red fixation cross and the thermode heated to the target temperature. Participants then withdrew their arm from the thermode and placed it beside the thermode. After the thermode returned to the baseline temperature participants rated the stimulus (“How intense was the stimulus?”) on a 0-100 visual analog scale (VAS) (0 = no sensation, 50 = minimally painful, 100 = unbearably painful), displayed for 6 s. The initial cursor position was randomized across trials. If participants did not withdraw during the stimulus, they were prompted to do so when the rating scale appeared, allowing the load cell to be re-tared during the subsequent 8-10 s intertrial interval. Two training trials, one at each temperature, preceded the first run.

#### Thermal rating task (fMRI experiment)

As part of the larger study, participants also completed a separate thermal rating task while inside the MRI scanner. From this task we obtained each participant’s pain ratings. Participants received the same fixed target temperatures as in the thermal withdrawal task (i.e. 46 °C and 48 °C), delivered to the left forearm and left lower leg with an MRI-compatible CHEPS thermode (identical model as used in the thermal withdrawal task). The baseline temperature was 32 °C. Target temperatures were reached and returned to baseline at a rate of 10°C/s (ramp durations 1.40 and 1.60 s) and held for an 8 s plateau. The task comprised four runs – two on the left forearm and two on the left lower leg (order counterbalanced across participants). Each run contained separate heat and cold blocks of 12 trials, with temperature order pseudorandomized within each block. Only the heat trials are analyzed here.

Each trial opened with a white fixation cross for 2s that turned red at stimulus onset. After a jittered 2-5 s delay, participants rated perceived intensity on the same VAS used in the thermal withdrawal task. Responses were entered with an MRI-compatible button box. The rating was followed by a 10-16 s interval containing different non-painful events (20 °C/40 °C thermal stimulation, a 2 s 1000 Hz tone, or a 2 s visual checkerboard).

#### Thermal pain calibration

Before the thermal withdrawal task, heat pain sensitivity was calibrated on the left forearm using a procedure adapted from Horing and Büchel [13]. Stimuli rose from a 32 °C baseline at 13 °C/s and were held for 10 s. The calibration consisted of two stages. First, the heat pain threshold was estimated from six painful/not-painful judgments using an adaptive maximum-likelihood procedure. Second, suprathreshold sensitivity was assessed with graded ratings: after two practice stimuli at and just above threshold, participants rated stimuli at temperatures predicted to elicit ratings of 10, 30 and 90, followed by stimuli that were updated online to target further rating levels (median 6 rated stimuli per participant, range 4–14), on the same VAS as in the other tasks.

### Signal processing

Signal processing was performed in MATLAB 2024a.

#### Accelerometer

All analyses used the acceleration magnitude (the Euclidean norm of the three axes, i.e., √(*x*² + *y*² + *z*²)). For each trial, a 12 s epoch was extracted from heat onset.

The magnitude signal was low-pass filtered with a 4^th^-order Butterworth filter (20 Hz cutoff). Movement onset was detected relative to a per-participant baseline, estimated from the first second of the stimulus pooled across trials: its location was the median, and its noise level the median absolute deviation (MAD), scaled to be consistent with the standard deviation. The absolute deviation from the baseline median was smoothed with a 15 ms moving average. In this signal, peaks exceeding 15 scaled MADs in height and 2.5 scaled MADs in prominence, at least 150 ms apart, were identified, and for each peak the movement onset was defined as the last preceding sample below 2 scaled MADs. The withdrawal was the first peak whose movement onset fell later than 1 s after stimulus onset. Withdrawal latency was defined as the interval between stimulus onset and this movement onset. Peak acceleration was defined as the maximum of the low-pass-filtered acceleration magnitude within 500 ms after movement onset, expressed in g (multiples of standard gravity, 9.81 m/s²). Because the magnitude includes the constant gravitational component, resting values are approximately 1 g and values above 1 reflect movement. Fig. 1C shows an example movement trace with the extracted variables annotated.

The peak criteria (first peak, minimum height 15 scaled MADs) were selected by agreement with the load cell, which registered loss of contact independently of the accelerometer.

#### Trial classifications and exclusions

Movement onsets were required to fall between 1 s and 11 s relative to stimulus onset. The upper bound was set to a common value for both temperatures, shortly before the end of the plateau (11.08 s at 46 °C and 11.23 s at 48 °C). Trials without a movement onset within this window were classified as endured, and their withdrawal latency was set to 11 s.

Of the 2,184 trials, 1,583 (72.5 %) yielded a measured withdrawal latency and 597 (27.3 %) were censored at 11 s as endured. Censoring occurred mostly at 46 °C (threshold: 196 of 543 trials; tolerance: 352 of 545) and rarely at 48 °C (threshold: 4 of 546; tolerance: 45 of 546; Table S1). Four trials (0.2 %) were excluded because movement onset could not be reliably determined.

### Statistical analyses

#### Effects of temperature and instruction

Effects of temperature and instruction on withdrawal latencies and peak acceleration were analyzed using separate linear mixed-effects models in R (version 4.6.1, lme4/lmerTest [2,16]). Fixed effects included temperature (46 °C, 48 °C), instruction (Threshold, Tolerance), and their interaction. Random effects were specified as maximal for the design [1], i.e., (1 + temperature ∗ instruction | participant). Where that fit was singular or failed to converge, it was reduced along a fixed sequence: first the random interaction slope was dropped, then the instruction slope, then the temperature slope. Withdrawal latency kept the maximal structure. For peak acceleration it was reduced to a by-participant intercept and slopes for temperature and instruction. Models were fitted by restricted maximum likelihood. Fixed effects were tested with Type III F-tests, using Satterthwaite’s approximation for the denominator degrees of freedom, with partial η² as the effect size. Condition means and temperature contrasts are reported as estimated marginal means (emmeans 2.0.4 [18]). Temperature effects are reported within each instruction (Holm-corrected across the two contrasts). Tests were performed two-sided with α = .05.

#### Association with the separate thermal rating task

To assess the association between the kinematic features and the by-participant mean ratings from the separate thermal rating task, we used Pearson correlations. Trial-level measures were aggregated to participant-level estimates within each participant × instruction × temperature cell, requiring at least three of the six trials in a cell; cells below that threshold were treated as missing. Withdrawal latency was the mean of the trial latencies in a cell, with endured trials entering at their censored value of 11 s. Peak acceleration was averaged only over trials in which a movement was detected. Each participant’s ratings of the 46 and 48 °C stimuli in the thermal rating task were averaged within temperature, across runs and stimulation sites. Correlations were computed separately for each measure, instruction and temperature. These eight tests were Holm-corrected as a single family. To quantify evidence for the absence of a correlation, we additionally computed Bayes factors (BF_01_; BayesFactor [22] default priors).

To test whether the associations differed between measures and between instructions, we compared the 48 °C correlations using Williams’ test for dependent, overlapping correlations [30]. Within each instruction, the withdrawal latency correlation was compared with the peak acceleration correlation. Within each measure, the threshold correlation was compared with the tolerance correlation. Each of the four tests was computed on participants with complete data for all three variables involved. These comparisons were not corrected for multiple comparisons.

Because a large proportion of trials at 46 °C ended without a withdrawal, restricting between-subject variance, we treat 48 °C as the main behavioral readout. The 46 °C correlations are reported alongside, and the corresponding figure is provided in the Supplementary Material.

#### Association with calibrated pain sensitivity

Two measures of pain sensitivity were derived from the calibration. The heat pain threshold was the temperature estimated by the adaptive procedure and was available for all 91 participants. For suprathreshold sensitivity, each participant’s calibration ratings were regressed linearly on stimulus temperature, and the temperatures predicted to elicit ratings of 50, 60, 70 and 80 were averaged (equivalent to the temperature predicted for a rating of 65). Participants with an outlying predicted temperature at any of these four ratings (Tukey’s 1.5 × IQR rule) were excluded from this measure, leaving 79 of 91 participants. For both measures, higher temperatures indicate lower pain sensitivity. Each measure was correlated (Pearson) with withdrawal latency and peak acceleration in each instruction × temperature cell, Holm-corrected as a separate family of eight per measure.

#### Withdrawal-task ratings

Ratings from the thermal withdrawal task were analyzed with a linear mixed model with the same fixed effects (temperature × instruction) and random-effects structure as the kinematic models. Tolerance and threshold ratings were compared at each temperature with simple contrasts from this model (Satterthwaite degrees of freedom). For visualization, ratings were also summarized separately for trials that ended in a withdrawal and trials that were endured. For the participant-level analyses, ratings were averaged within each participant × instruction × temperature cell, using the same ≥ 3-of-6-trial rule as for the kinematic measures. At 48 °C, these mean ratings were correlated (Pearson) with the ratings from the thermal rating task and with both kinematic measures. We also computed partial correlations between each kinematic measure and the thermal-rating-task rating, controlling for the withdrawal-task rating. These analyses were not corrected for multiple comparisons.

#### Reliability

Reliability of each measure was estimated as permutation-based split-half internal consistency. Within each temperature × instruction condition, each participant’s trials were randomly split into two halves. Participant means were computed for each half and correlated across participants, and the correlation was corrected to full test length with the Spearman-Brown formula. Participants with fewer than four valid trials in a condition were left out. This was repeated over 5,000 random splits, and the median and 95% interval of the resulting distribution are reported. Reliability of the thermal-rating-task rating was estimated in the same way.

#### Robustness analyses

We assessed the robustness of the main results to the treatment of endured trials and to other analytic choices.

Entering endured trials at 11 s treats a latency of at least 11 s as an observed value, which compresses the latency distribution, most strongly at 46 °C. We therefore refitted the latency model as a censored mixed model, with endured trials right-censored at 11 s and the same fixed and random effects as the linear mixed model. A Bayesian implementation was used because it allowed right-censoring to be combined with the maximal random-effects structure. The model was fitted in brms 2.23.0 [3] with a Gaussian likelihood and default priors (flat on the fixed effects; 4 chains of 3,000 iterations, including 1,000 warm-up; latency expressed relative to the 11 s ceiling and all chains initialized at zero; all R^ < 1.01). Its estimates refer to latent, uncensored withdrawal latencies. To separate the effect of censoring from differences between estimation frameworks, the same model was also fitted without censoring in brms. We report posterior means, 95% equal-tailed credible intervals and the probability of direction (pd).

To test whether the latency–rating association at 46 °C reflects censoring, we re-estimated it as a censored regression of trial-level latency on the thermal-rating-task rating, with endured trials right-censored at 11 s, fitted in brms with the same settings as the censored latency model. The model estimated a separate rating slope and residual SD for each instruction × temperature cell, with uncorrelated by-participant random intercepts per cell. Slopes are expressed in seconds per 10 VAS points and reported per cell.

Finally, we recomputed the associations with the thermal rating task as Spearman rank correlations, recomputed the latency correlations after discarding endured trials instead of entering them at 11 s, and restricted the thermal-rating-task rating to stimuli applied to the arm or to the leg.

## Results

### Effects of temperature and instruction on withdrawal features

As expected, withdrawal latencies decreased with stimulus temperature (*F*(1, 89.99) = 1045.00, *p* = 2.57 × 10^−51^, η^2^_p_ = .92; Fig. 2A). Participants withdrew 3.65 s earlier at 48 °C than at 46 °C under the threshold instruction and 3.95 s earlier under the tolerance instruction (Fig. 2C). This effect held when endured trials were modeled as censored instead of entered as 11 s (Table S2). Peak acceleration increased with stimulus temperature (*F*(1, 90.57) = 25.46, *p* = 2.31 × 10^−6^, η^2^_p_ = .22; Fig. 2B), by 0.075 g under threshold and 0.119 g under tolerance (Fig. 2D; Table S3b).

**Fig. 2.**
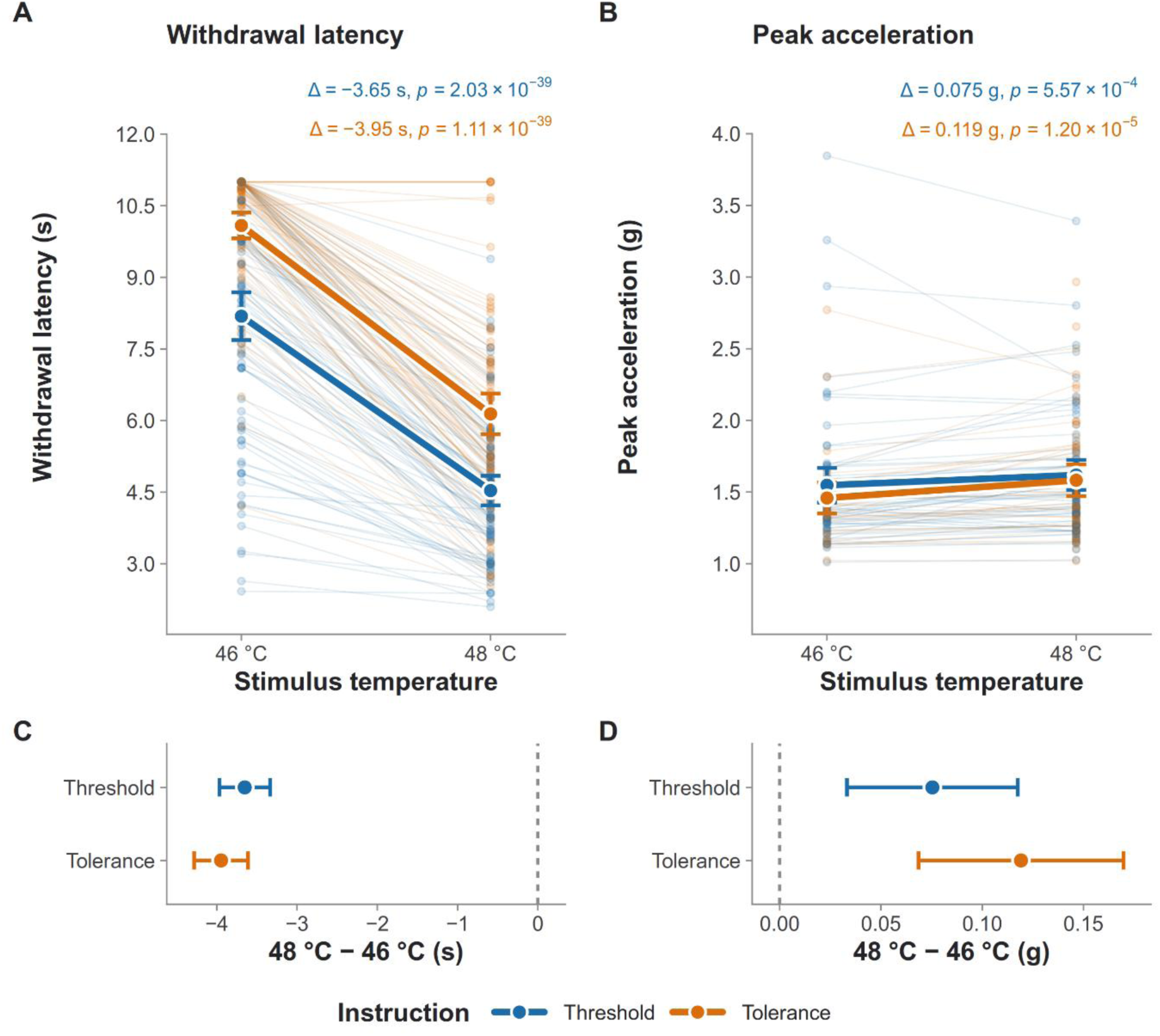
Withdrawal features across stimulus intensities and instructions. Withdrawal latency **(A)** and peak acceleration **(B)** at 46 °C and 48 °C under pain threshold (blue) and tolerance (orange) instructions. As stimulus temperature increases, withdrawal latency decreases while peak acceleration (withdrawal vigor) increases. Faint points and connected lines represent individual participant means. Heavy dots represent group means with 95% confidence intervals. Annotations display the estimated difference (Δ, 48 °C − 46 °C) and the associated *p*-values, derived from contrasts of estimated marginal means in linear mixed models (Holm-adjusted). **(C, D)** Estimated temperature effects (48 °C − 46 °C) on withdrawal latency (C) and peak acceleration (D) under each instruction, with 95% confidence intervals; the dashed line marks no difference. Trials without a motor withdrawal are capped at the 11.0 s maximum for latency in (A, C) and are missing from the peak acceleration panels (B, D). The threshold run always preceded the tolerance run.

Withdrawal latencies were longer in the tolerance than in the threshold run (*F*(1, 90.05) = 117.67, *p* = 5.01 × 10^-18^), whereas peak acceleration did not differ significantly between runs (*F*(1, 93.79) = 3.12, *p* = .081; Table S3). Because the threshold run always preceded the tolerance run, these differences cannot be separated from order effects and are not interpreted further.

Both withdrawal latency and peak acceleration were reliably measured. All split-half reliabilities exceeded .90 at 48 °C (Fig. S1).

Withdrawal latency and peak acceleration shared little between-subject variance and were not significantly correlated in any condition (all |*r*| ≤ .23; Table S4).

### Inter-individual associations with pain sensitivity

Participants who withdrew earlier rated the heat stimuli as more painful in the thermal rating task (Fig. 3A). At 48 °C, withdrawal latency correlated with mean pain ratings at *r* = −.468 (95% CI [−.614, −.290], *n* = 91, *p_adj_* = 2.08 × 10^−5^) under the threshold instruction and at *r* = −.501 (95% CI [−.641, −.330], *n* = 91, *p_adj_* = 3.29 × 10^−6^) under the tolerance instruction. Peak acceleration showed no such relationship under either instruction (threshold *r* = −.013 [−.220, .194], *n* = 90; tolerance *r* = −.050 [−.262, .166], *n* = 84, both *p_adj_* = 1.00; Fig. 3B), with Bayes factors providing moderate evidence for the null (BF_01_ = 4.1 and 3.6).

**Fig. 3.**
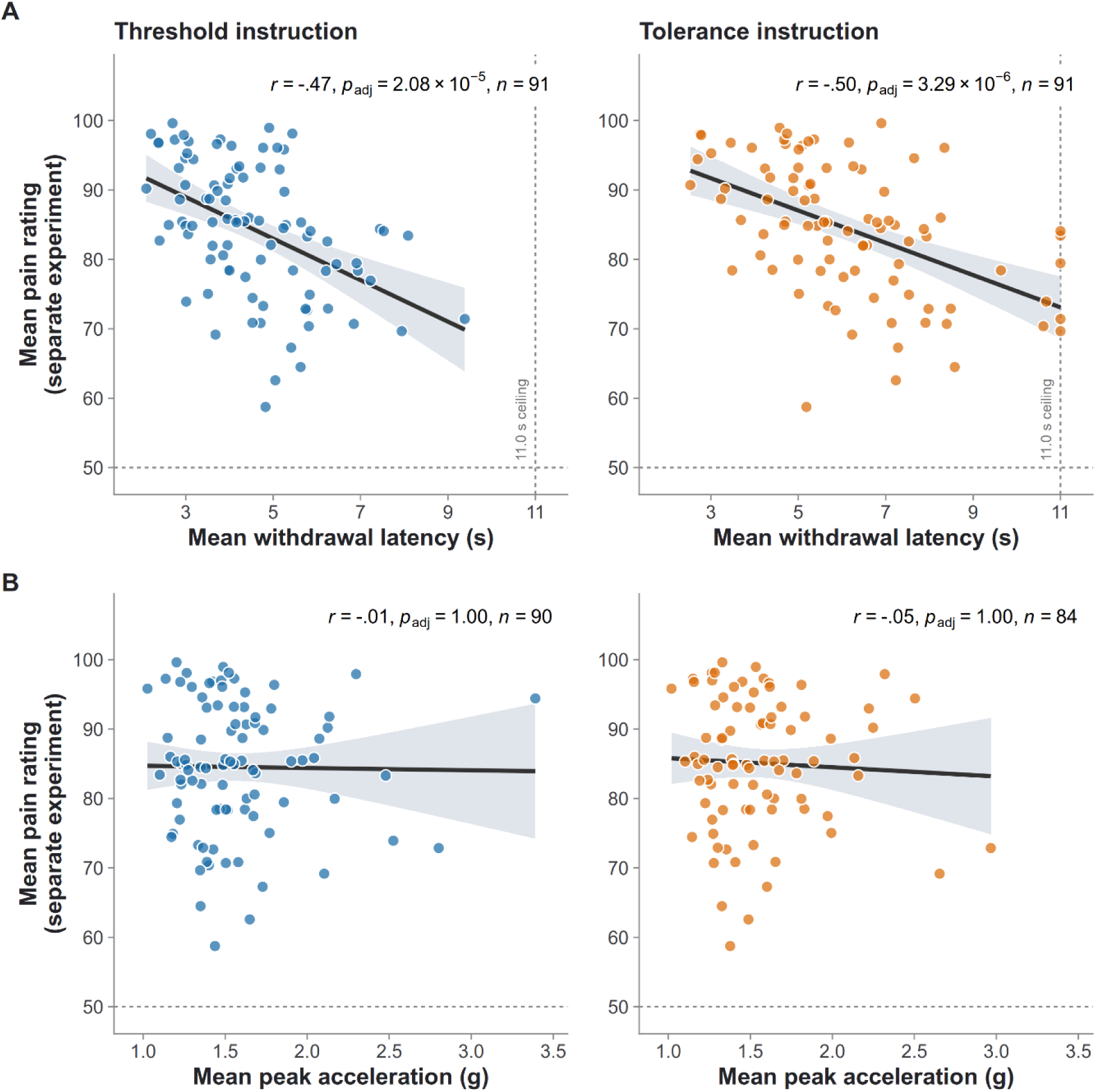
Association between withdrawal features and separate pain ratings. **(A)** Mean withdrawal latency and **(B)** mean peak acceleration in response to the 48 °C stimulus, and mean pain ratings derived from a separate thermal rating task (fMRI experiment; 0–100 scale, 50 = minimally painful). Responses are shown for the pain threshold (left, blue) and tolerance (right, orange) instructions. Each point represents a single participant. Solid lines indicate ordinary least-squares regression fits, with shaded areas representing 95% confidence intervals. Annotations provide Pearson’s correlation (*r*), p-values (Holm-adjusted within the family of eight kinematic correlations across both temperatures), and sample size (*n*). Withdrawal latency correlates negatively with separate pain ratings across both instructions, whereas peak acceleration shows no relationship. The 48 °C condition is visualized here because the higher stimulus intensity elicited fewer non-withdrawal trials, increasing the number of measurable withdrawal responses; the associations for the 46 °C condition are provided in Fig. S2. In (A), participants who endured the full stimulus without withdrawing are retained and censored at the 11.0 s observation ceiling (dashed vertical line). Because acceleration amplitude cannot be extracted without a withdrawal movement, these non-withdrawal trials are excluded from (B).

The association with the separate ratings was significantly stronger for withdrawal latency than for peak acceleration under both the threshold (Williams’ *t*(87) = −2.90, *p* = .005) and the tolerance instruction (*t*(81) = −2.74, *p* = .008). Neither association differed between instructions (withdrawal latency: *t*(88) = 0.47, *p* = .641; peak acceleration: *t*(81) = 0.96, *p* = .341).

At 46 °C, the association between withdrawal latency and the separate pain ratings was weaker but held under both instructions (Fig. S2; threshold *r* = −.364, 95% CI [−.530, −.170], p_adj_ = .002; tolerance *r* = −.318, [−.492, −.120], p_adj_ = .010). The weaker association might reflect the stimulus ceiling: 36 % and 65 % of trials at 46 °C ended without a withdrawal, compared with 1 % and 8 % at 48 °C (Table S1, Fig. S3). A censored regression, which treats these trials as right-censored, showed a reliable association in all four conditions (Table S5).

The reported associations did not depend on analytic choices. At 48° C, the association between withdrawal latency and the separate pain ratings was robust to correlation method (Spearman *ρ* = −.50 and −.54), to censoring treatment (censored regression; endured trials dropped, *r* = −.45 and −.47), and to how the separate rating was constructed (all variants |*r*| ≥ .40), whereas peak acceleration showed no association under any variant (all |*r*| ≤ .10; Table S5).

Withdrawal latency was also related to both measures from the pre-task calibration (higher temperatures indicate lower sensitivity; Fig. 4; Fig. S4; Fig. S5). Participants with lower heat pain thresholds withdrew earlier at both temperatures and under both instructions (*r* = .38 to .43, *n* = 91, all *p_adj_* ≤ .002). Likewise, participants with higher suprathreshold sensitivity withdrew earlier in all conditions (*r* = .37 to .55, *n* = 79, all *p_adj_* ≤ .005). In contrast, peak acceleration showed no significant association with either calibration measure in any condition (heat pain threshold: all *p_adj_* ≥ .757; suprathreshold sensitivity: all *p_adj_* ≥ .535). Fig. 4 summarizes the associations of both withdrawal features with all three pain sensitivity measures.

**Fig. 4.**
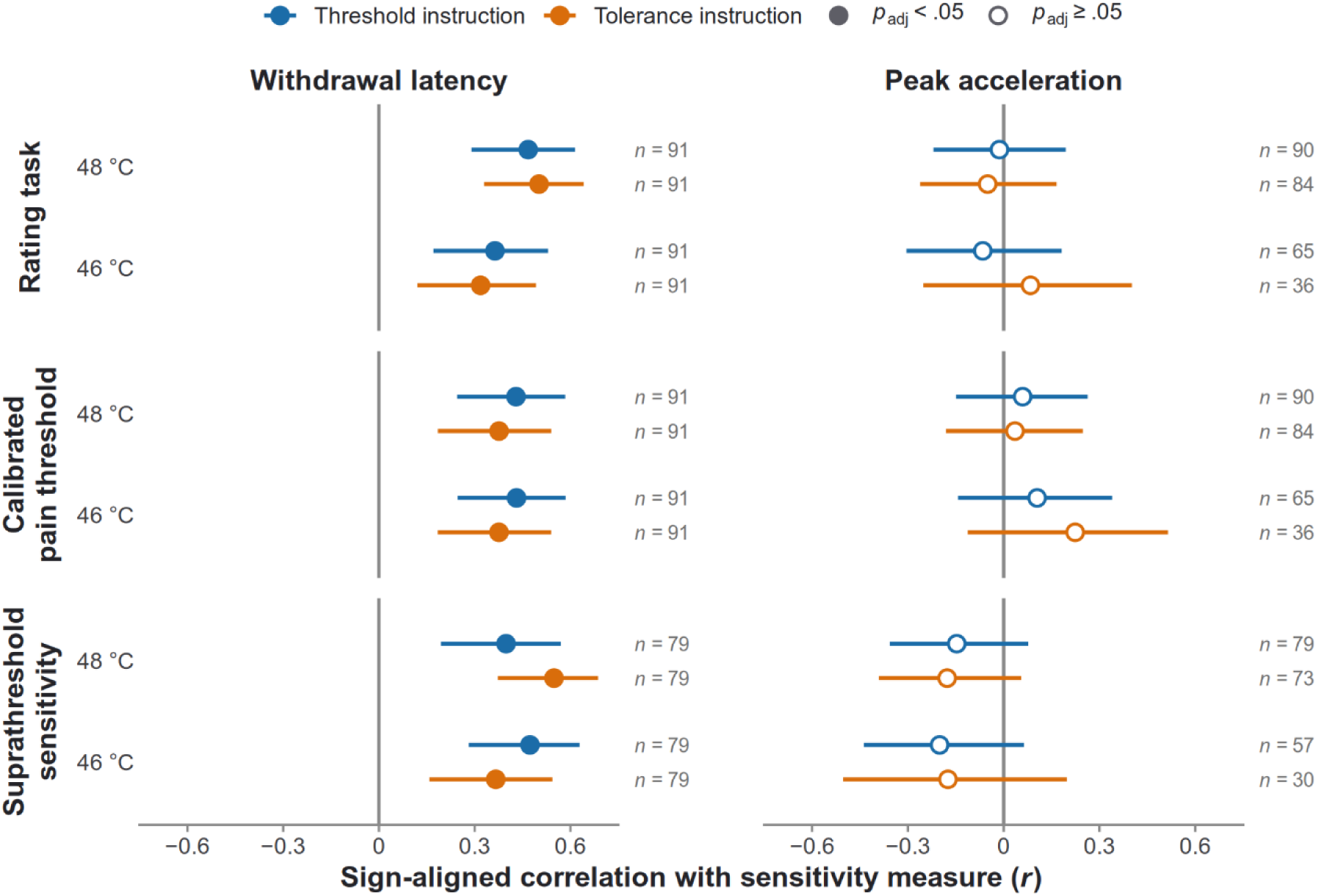
Associations of withdrawal features with different measures of pain sensitivity. Pearson correlations of withdrawal latency (left) and peak acceleration (right) with mean pain ratings from the separate thermal rating task, the heat pain threshold from the pre-task calibration, and suprathreshold sensitivity from the pre-task calibration, at 48 °C and 46 °C under the pain threshold (blue) and tolerance (orange) instructions. Correlations were sign-aligned so that positive values indicate that more pain-sensitive participants withdrew earlier (latency) or with higher peak acceleration. Error bars represent 95% confidence intervals. Filled symbols indicate *p* < .05 after Holm correction within each sensitivity measure’s family of eight tests; open symbols indicate non-significant correlations. Sample sizes (*n*) are given for each correlation; because peak acceleration requires a withdrawal response, *n* is lower where many trials were endured.

### Thermal withdrawal task ratings

Ratings collected during the withdrawal task were consistent with the instructed criteria (Fig. S6). Ratings were higher under the tolerance than under the threshold instruction at both 46 °C (*M* = 59.5 vs. 53.1; difference = 6.4, 95% CI [4.4, 8.5], *t*(90) = 6.26, *p* = 1.29 × 10^−8^) and 48 °C (78.4 vs. 65.5; difference = 12.9, 95% CI [10.5, 15.3], *t*(90) = 10.83, *p* = 5.46 × 10^−18^), and this difference was larger at 48 °C (instruction × temperature interaction: *b* = 6.5, *t*(90) = 5.89, *p* = 6.44 × 10^−8^).

At 48 °C, withdrawal-task ratings correlated with the mean pain ratings from the separate thermal rating task under the threshold instruction (*r* = .35, *p* = 7.88 × 10^−4^) and under the tolerance instruction (*r* = .24, *p* = .021). They did not correlate with withdrawal latency (threshold: *r* = −.03, *p* = .810; tolerance: *r* = .05, *p* = .658) or peak acceleration (*r* = −.07, *p* = .537; *r* = .03, *p* = .759). Controlling for the withdrawal-task ratings left the association between latency and separate ratings unchanged (partial *r* = −.49, 95% CI [−.63, −.31], *p* = 1.00 × 10^−6^ and partial *r* = −.53, [−.66, −.36], *p* = 8.21 × 10^−8^; zero-order *r* = −.47 and −.50), and the peak-acceleration association remained null (partial *r* = .01, *p* = .926 and −.06, *p* = .586).

At 46 °C, where trials ending without a withdrawal received lower ratings (Fig. S6), withdrawal-task ratings correlated with withdrawal latency (threshold: *r* = −.41, *p* = 4.71 × 10^−5^; tolerance: *r* = −.29, *p* = .005). Controlling for them, the association between latency and separate ratings remained (partial *r* = −.25, *p* = .015 and −.24, *p* = .022; Table S5).

## Discussion

Using a human hot plate analog, we recorded temporal and kinematic features of withdrawal responses to fixed thermal noxious stimuli. Intra-individually, withdrawal latency decreased and peak acceleration increased with stimulus intensity. Crucially, withdrawal latency also tracked inter-individual differences in pain sensitivity across three separate measures (pain ratings in an independent task, the heat pain threshold and suprathreshold sensitivity). By quantifying movement kinematics, we could further test how participants withdrew. Peak acceleration showed no relationship with pain ratings despite high measurement reliability, indicating that inter-individual differences in pain sensitivity are reflected in the timepoint at which participants withdraw, but not in how vigorously they do so.

The intra-individual effects of temperature confirm that nocifensive withdrawal behavior scales with stimulus intensity. Consistent with previous findings in both animal models [9,11] and a human thermal withdrawal analog [10], higher temperatures reliably elicited shorter withdrawal latencies. Crucially, in our paradigm, these scaling effects extend beyond the timing of the movement to the intensity of its execution. Withdrawal vigor, indexed by peak acceleration, increased with stimulus intensity. This aligns with electromyography evidence in both animal models [5] and humans [4] showing greater muscle recruitment for more intense stimuli, and with the observation that noxious stimulus intensity can facilitate generalized motor behavior, as indexed by shorter pain-related reaction times [12,19,25,28]. Thus, participants withdraw not only earlier but with greater withdrawal vigor at higher temperatures. This demonstrates that wrist-mounted accelerometry provides a sensitive, continuous measure of this intensity-dependent motor response, capturing graded changes in withdrawal vigor that complement traditional latency-based measures.

While both temporal and kinematic features scaled intra-individually with stimulus intensity, only withdrawal latency captured inter-individual differences in pain sensitivity. Participants who withdrew earlier consistently reported higher pain ratings in a separate temperature-matched task and had lower heat pain thresholds and higher suprathreshold sensitivity in a calibration performed before the withdrawal task. As our paradigm uses fixed-temperature stimuli, the withdrawal latency reflects how quickly the nociceptive input reaches the participant’s subjective criterion – whether instructed to withdraw at their pain threshold or tolerance. Individuals with greater pain sensitivity reach these internal criteria faster, and, accordingly, rate the same stimuli higher in a separate task. In contrast, despite excellent measurement reliability, the data favored the absence of a relationship between peak acceleration (i.e., vigor) and pain ratings. This divergence suggests that the link between pain sensitivity and nocifensive behavior is reflected in the evaluative or decisional stage – captured by withdrawal latency – rather than in motor execution. Once the decision threshold is reached, the physical withdrawal execution appears to be driven by the stimulus itself, scaling with temperature but not with inter-individual differences in pain sensitivity. Consequently, withdrawal latency serves as a behavioral readout of pain sensitivity, whereas measuring withdrawal vigor alongside helps to disentangle the subjective decision to withdraw from stimulus-driven motor execution.

An alternative explanation is that differences in withdrawal latency reflect variations in subjective pain intensity at withdrawal. However, by design, the withdrawal instructions anchor the moment of withdrawal to a specific subjective criterion, constraining variability of the momentary pain experience. Accordingly, at 48 °C, withdrawal-task ratings were consistent with the instructed criteria but showed no relationship with withdrawal latency, and controlling for them left the association between withdrawal latency and separate pain ratings unchanged. At 46 °C, where trials ending without a withdrawal were rated lower, the two were correlated, but the association with separate pain ratings remained after controlling for them. Withdrawal latency is therefore not merely a motor proxy for concurrent pain perception. Rather, it captures a facet of pain sensitivity, manifesting across contexts as both earlier behavioral withdrawal and higher subjective pain ratings. This pattern of stable inter-individual differences is consistent with the rodent literature, where thermal withdrawal latency captures reliable inter-individual differences [26] and demonstrates heritability [21].

Our paradigm addresses a fundamental limitation in human pain research: the artificial suppression of movement. As noted previously [6,25], traditional human pain assessments, particularly in neuroimaging paradigms, typically require participants to remain still. This methodological constraint conflicts with one core evolutionary function of pain, which is to drive protective withdrawal [29]. By allowing participants to execute a self-paced withdrawal response, our hot plate analog provides a closer approximation of naturalistic pain behavior than traditional static paradigms. Such an action-oriented framework facilitates a more direct comparison with preclinical assays.

By establishing a shared behavioral readout – withdrawal latency – our findings offer a translational link between preclinical assays and human pain studies. Although rodent thermal withdrawal tasks are a cornerstone of preclinical assays, translating these behaviors across species is challenging [20,24]. Correlating withdrawal latency with subjective pain is untestable in rodents, where pain can only be inferred from behavior itself [8]. Addressing this, our paradigm retains the defining features of these preclinical tests – using a self-paced behavioral escape from a noxious stimulus as the primary endpoint – but applies it in humans, where it can be validated against subjective experience. Furthermore, this framework allows us to decouple the evaluative decision to withdraw from the actual motor execution. While withdrawal latency reliably tracks pain sensitivity, the physical vigor of the response does not. For interpreting preclinical data, this suggests that the magnitude of a motor response may not reliably reflect inter-individual differences in pain sensitivity, pointing to latency as the more robust cross-species index of pain.

Withdrawal latency may also allow pain sensitivity to be tested where self-report is limited. Kamper-Fuhrmann and colleagues [14] proposed hand withdrawal as a simpler and safer alternative to standard psychophysics for young children or cognitively impaired individuals. Our findings add that, at a fixed temperature, withdrawal latency tracks inter-individual differences in pain sensitivity, supporting its use as a behavioral form of sensory testing that requires no rating. The paradigm could further be adapted to free withdrawal without a predefined criterion, potentially extending sensory testing to individuals who cannot follow instructions.

At 46 °C, many participants endured the full stimulus without withdrawing, which compressed latencies at the ceiling and likely weakened the correlations with pain ratings. However, withdrawal latency was associated with pain ratings at this temperature under both instructions, including when endured trials were modeled as censored. At 48 °C, where almost all trials ended in a withdrawal, the association was strongest.

In summary, withdrawal latency in a human hot plate analog captured inter-individual differences in pain sensitivity across three separate measures, providing a behavioral, non-verbal complement to traditional self-report. Furthermore, incorporating movement kinematics isolates the cognitive decision to withdraw from the physical motor execution. Crucially, by sharing its primary endpoint with the rodent hot plate test while allowing validation against subjective report, this paradigm offers a framework for translating between human and animal models of pain.

## Data and code availability

The dataset underlying all analyses and figures can be accessed at [GIN repository, https://gin.g-node.org/lrsktr/hotplate_data] released under a CC BY-NC 4.0 license. Processing scripts, statistical analysis code, and figure-generation code are provided at [GIN repository, https://gin.g-node.org/lrsktr/hotplate_code] under the MIT license.

## Acknowledgements

We thank Helene Rittmeister for assistance with data collection and Mona Wiegers for assistance with designing the apparatus schematic.

This work was supported by the European Research Council (grant number 883892, PainPersist) and the German Federal Ministry of Education and Research (BMBF).

The authors used a large language model (Claude Opus 5.5, Anthropic) for coding assistance and English language editing during manuscript preparation.

The authors have no conflicts of interest to declare.

