## Supplementary Material for "Withdrawal latency in a human hot plate analog tracks inter-individual differences in pain sensitivity"

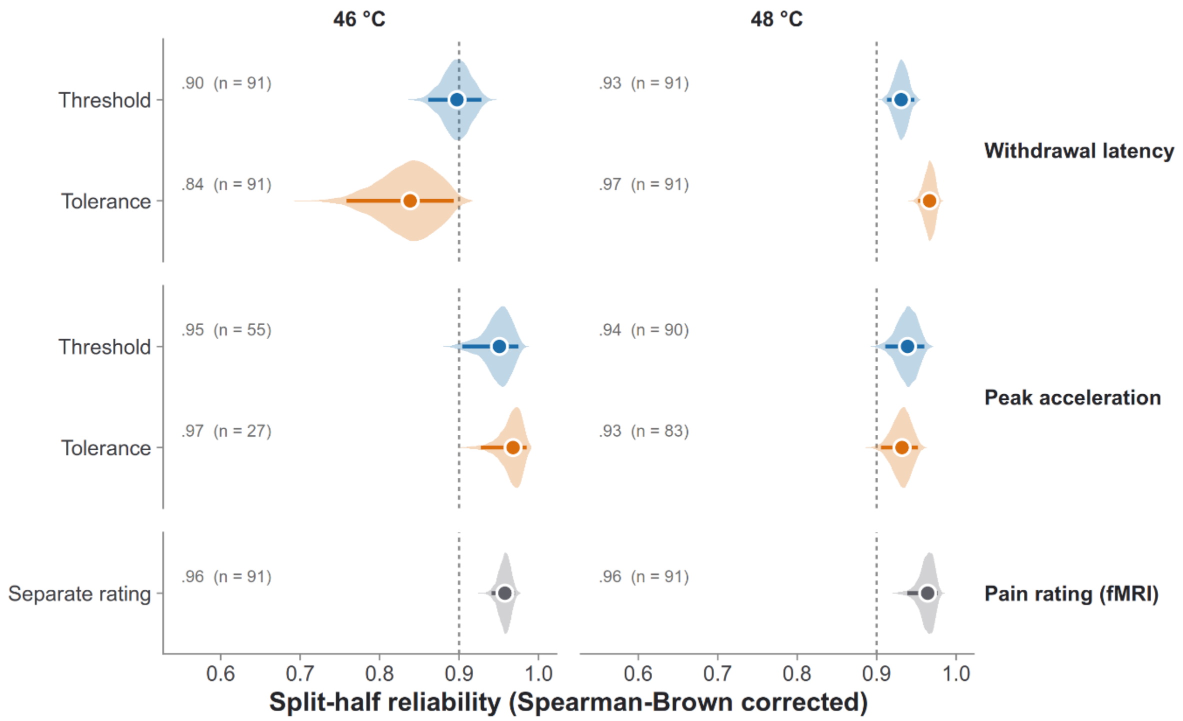


**Supplementary Figure S1. Split-half reliability distributions for withdrawal features and separate pain ratings.** Violin plots illustrate the distribution of split-half reliabilities for withdrawal latency (top), peak acceleration (middle), and pain ratings from the separate thermal rating task in response to 46 °C (left) and 48 °C (right) stimuli. Distributions are separated by the pain threshold (blue) and tolerance (orange) instructions. A vertical dashed line at 0.9 is provided as a reference for excellent reliability. To assess internal consistency, each participant's trials within a specific condition were randomly partitioned into two halves; participant means were computed on each half and correlated across participants. This permutation process was repeated over 5,000 random splits per cell, with a Spearman-Brown correction applied to the resulting correlations. The central white marker and solid horizontal bar represent the median and 95% interval of the distribution, respectively. Text annotations denote the median reliability and the sample size (n) of participants who had at least four valid trials in that cell. For withdrawal latency at 46 °C, trials where participants endured the stimulus are retained and censored at 11.0 s. For peak acceleration at 46 °C, the sample is restricted to participants who physically withdrew; consequently, the sample size is notably reduced under the tolerance instruction (n = 27) because most participants endured the stimulus.

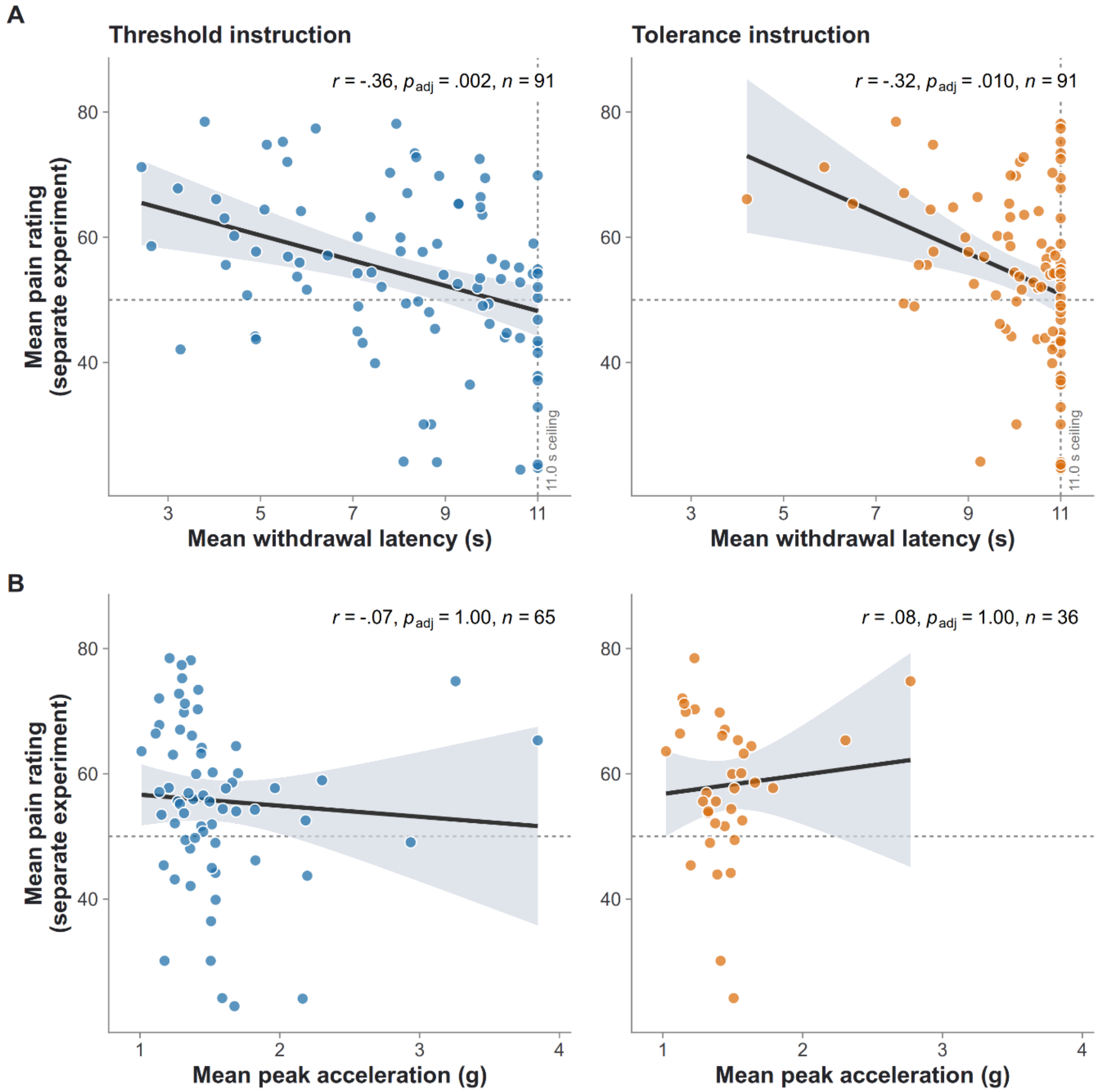


**Supplementary Figure S2. Association between withdrawal features and separate pain ratings at 46 °C**. **(A)** Mean withdrawal latency and **(B)** mean peak acceleration in response to the 46 °C stimulus, and mean pain ratings derived from a separate thermal rating task (fMRI experiment; 0–100 scale, 50 = minimally painful). The horizontal dashed line indicates the visual analog scale anchor for "minimally painful" (50). Responses are shown for the pain threshold (left, blue) and tolerance (right, orange) instructions. Each point represents a single participant. Solid lines indicate ordinary least-squares regression fits, with shaded areas representing 95% confidence intervals. Annotations provide Pearson's correlation (*r*), *p*-values (Holm-adjusted within the family of eight kinematic correlations across both temperatures), and sample size (*n*). Withdrawal latency correlates negatively with separate pain ratings, whereas peak acceleration shows no relationship. In (A), participants who endured the full stimulus without withdrawing are retained and censored at the 11.0 s observation ceiling (dashed vertical line). Censoring-aware estimates are detailed in Supplementary Table S5. Because acceleration amplitude cannot be extracted without a withdrawal movement, these non-withdrawal trials are excluded from (B). Consequently, panel (B) only includes participants with at least three valid withdrawals (n = 65 under threshold, n = 36 under tolerance). Under the tolerance instruction, where the majority of trials were endured, this exclusion yields a subsample inherently selected for higher pain sensitivity.

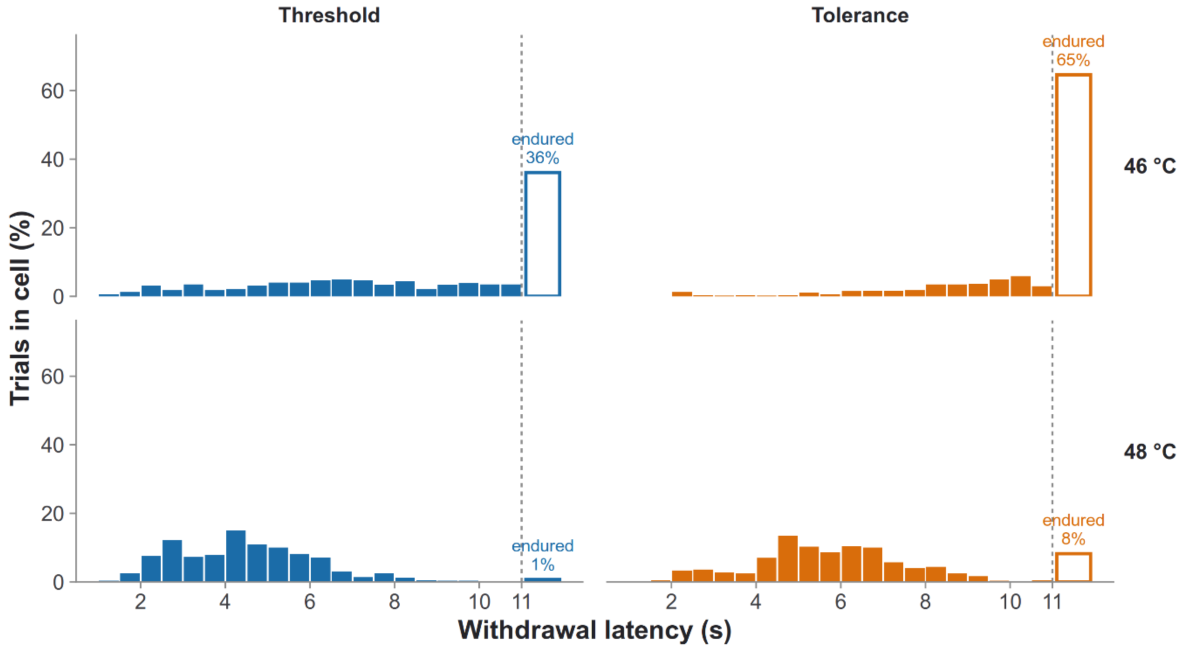


**Supplementary Figure S3. Distributions of withdrawal latencies and observation ceiling effects.** Histograms displaying the distribution of withdrawal latencies across 46 °C (top row) and 48 °C (bottom row) stimuli under the pain threshold (left, blue) and tolerance (right, orange) instructions. Solid bars represent measured withdrawal latencies grouped into 0.5 s bins, expressed as a percentage of trials within that condition. Open bars situated at the vertical dashed line represent trials in which the participant endured the full stimulus and never withdrew. Annotations above the open bars denote the exact percentage of endured trials per condition. The ceiling effect at 46 °C (up to 65% under the tolerance instruction) illustrates the restricted variance in this condition.

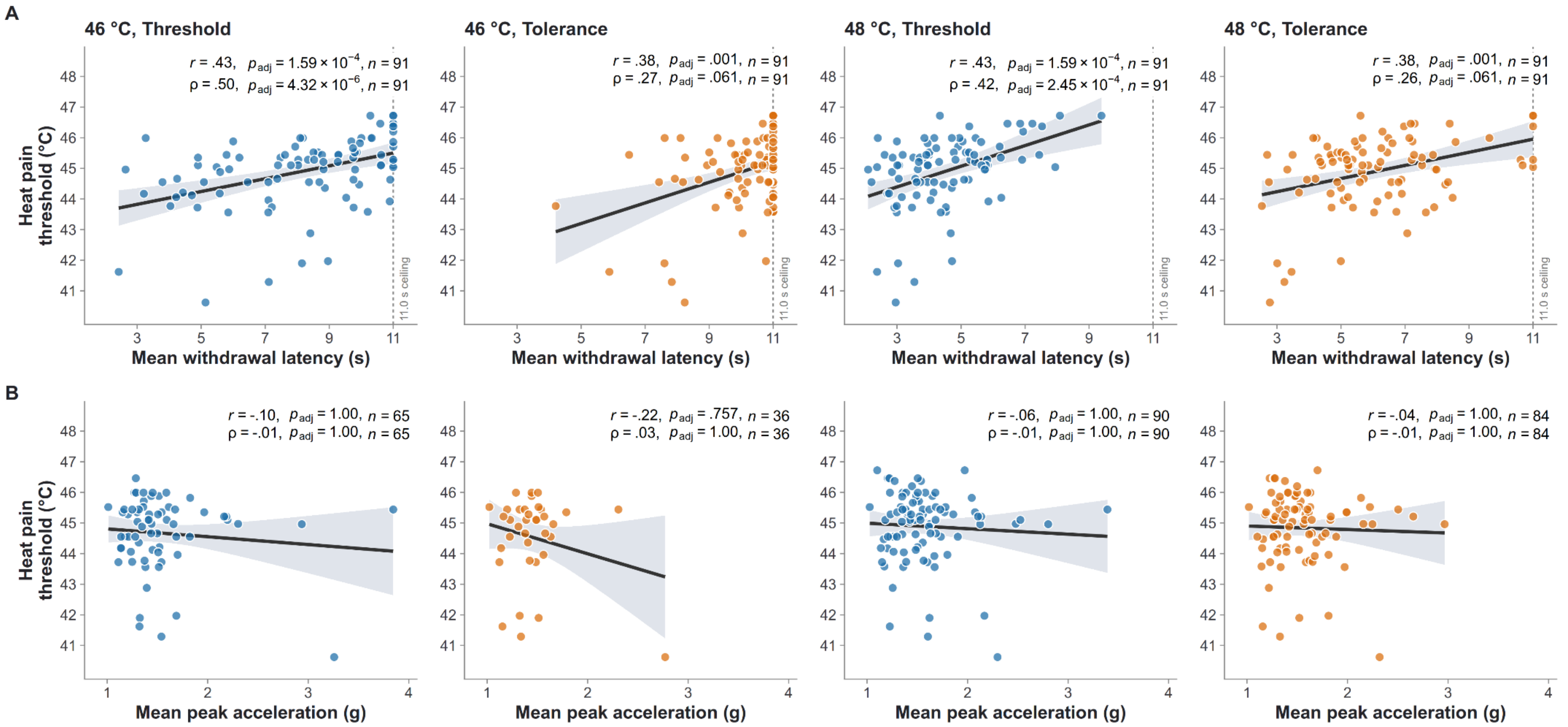


**Supplementary Figure S4. Withdrawal features and heat pain threshold.** Mean withdrawal latency (A) and mean peak acceleration (B) plotted against the heat pain threshold for each temperature, under the pain threshold (blue) and tolerance (orange) instructions. The heat pain threshold (y axis) was estimated with an adaptive maximum-likelihood procedure in the pre-task calibration; higher values indicate lower sensitivity. Each point represents one participant. The threshold was available for all 91 participants, and no exclusions were applied. Solid lines show ordinary least-squares fits with 95% confidence intervals. Annotations give Pearson's $r$, $p$-values (Holm-adjusted within the eight heat pain threshold correlations) and $n$. Additional annotations give Spearman's ρ and $p$-values (Holm-adjusted within the eight Spearman correlations). In (A), endured trials are entered at the 11.0 s observation ceiling (dashed vertical line). Because peak acceleration can only be extracted from trials with a withdrawal, the panels in (B) include fewer participants.

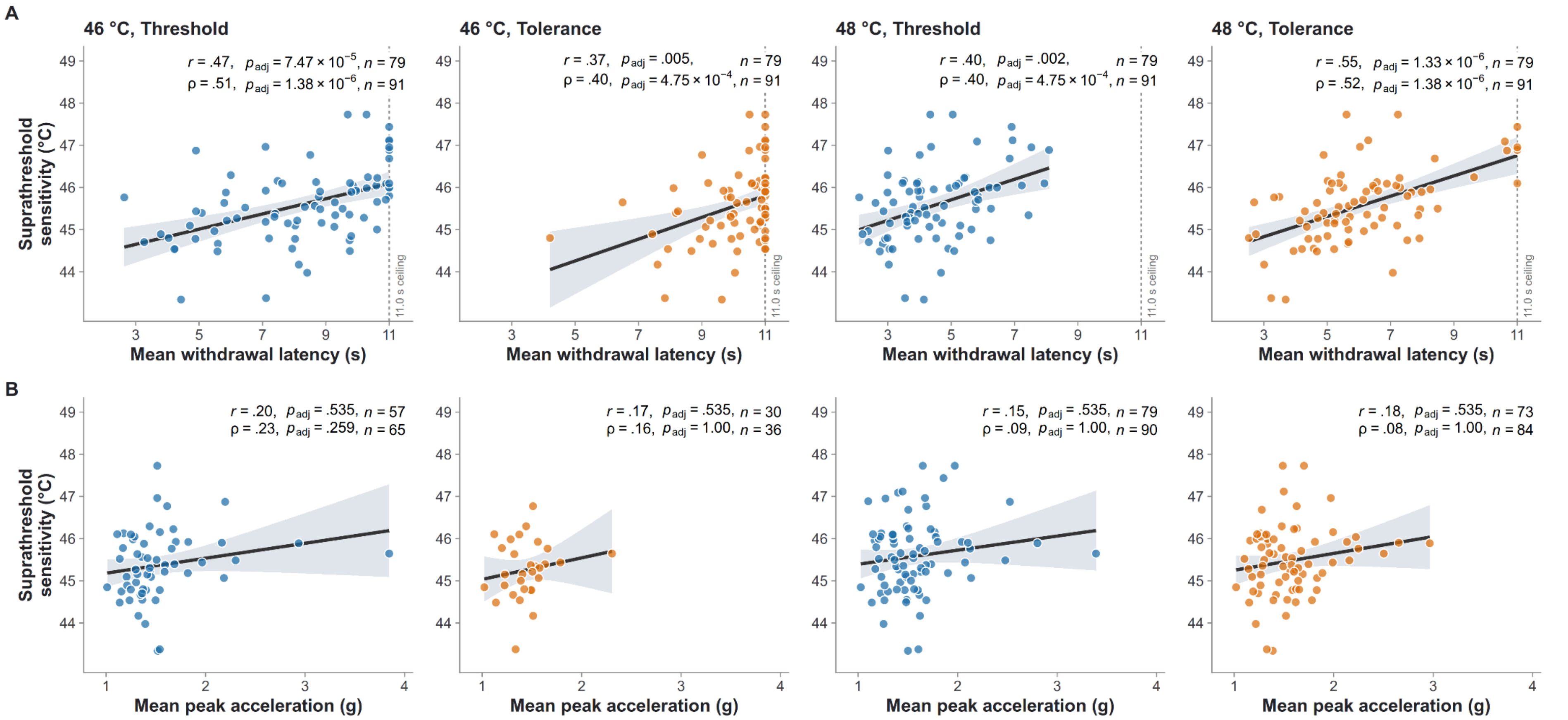


**Supplementary Figure S5. Withdrawal features and suprathreshold sensitivity.** Mean withdrawal latency (A) and mean peak acceleration (B) plotted against suprathreshold sensitivity for each temperature, under the pain threshold (blue) and tolerance (orange) instructions. Suprathreshold sensitivity (y axis) is the mean of the temperatures predicted to elicit ratings of 50, 60, 70 and 80 in the pre-task calibration (equivalent to the temperature predicted for a rating of 65); higher values indicate lower sensitivity. Each point represents one participant. Of the 91 participants, 12 were excluded (Tukey's 1.5 × IQR criterion, applied at each of the four target ratings), leaving *n* = 79. Excluded participants are not shown. Solid lines show ordinary least-squares fits with 95% confidence intervals. Annotations give Pearson's *r*, *p*-values (Holm-adjusted within the eight suprathreshold sensitivity correlations) and *n*. Additional annotations give Spearman's ρ and *p*-values computed on all 91 participants without the outlier exclusion (Holm-adjusted within the eight Spearman correlations). In (A), endured trials are entered at the 11.0

s observation ceiling (dashed vertical line). Because peak acceleration can only be extracted from trials with a withdrawal, the panels in (B) include fewer participants. At 46 °C under the tolerance instruction, these are mainly participants who withdrew and who therefore tend to be more pain sensitive.

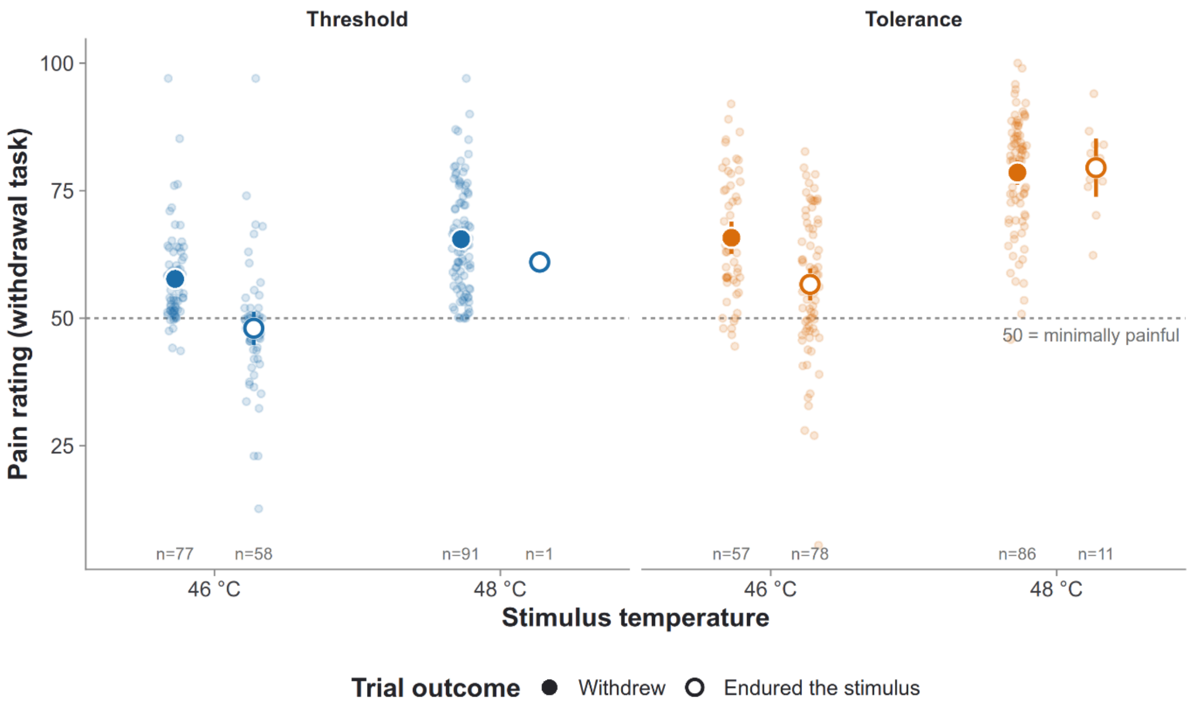


**Supplementary Figure S6. Withdrawal-task pain ratings for endured and withdrawn trials.** Mean pain intensity ratings (0–100 VAS) obtained during the thermal withdrawal task for 46 °C and 48 °C stimuli under pain threshold (left, blue) and tolerance (right, orange) instructions. Faint, smaller points represent individual participant means. Large circles represent group means, with error bars indicating 95% confidence intervals. Solid circles denote trials in which the participant executed a motor withdrawal; open circles denote trials in which the participant endured the full 10 s stimulus plateau. Sample sizes (*n*) for each subgroup are provided at the bottom of the plot. Subgroup sample sizes are not mutually exclusive, as participants who withdrew on some trials and endured on others contribute to both groups. The horizontal dashed line indicates the visual analog scale anchor for "minimally painful" (50).

**Supplementary Table S1.** Censoring of withdrawal latency by condition

| Instruction | Temperature | Total trials | Withdrew (latency measured) | Endured (censored at 11.0 s) | Excluded (no detectable onset) | % endured of analyzed trials | Participants never withdrawing |
|---|---|---|---|---|---|---|---|
| Threshold | 46 °C | 546 | 347 | 196 | 3 | 36.1 | 14 / 91 |
| | 48 °C | 546 | 542 | 4 | 0 | 0.7 | 0 / 91 |
| Tolerance | 46 °C | 546 | 193 | 352 | 1 | 64.6 | 34 / 91 |
| | 48 °C | 546 | 501 | 45 | 0 | 8.2 | 5 / 91 |

*Note.* Total = withdrew + endured + excluded. Endured: no withdrawal within the 11.0 s observation window; latency is censored there. Excluded: no withdrawal onset could be determined from the accelerometer, so the trial has no latency. Analyzed trials = withdrew + endured. Participants never withdrawing: every analyzed trial in the condition was endured, so the participant's mean latency equals the ceiling.

**Supplementary Table S2.** Withdrawal latency models accounting for right-censoring at 11.0 s

| Model | Temperature (48 − 46 °C) | Instruction (tolerance − threshold) | Temperature × instruction |
|---|---|---|---|
| Linear mixed model (primary; endured trials = 11.0 s) | −3.80 [−4.03, −3.57], $p = 2.57 \times 10^{-51}$ | 1.75 [1.44, 2.07], $p = 5.01 \times 10^{-18}$ | −0.29 [−0.74, 0.15], $p = .201$ |
| Bayesian mixed model, uncensored (endured trials = 11.0 s) | −3.80 [−4.04, −3.55], *pd* = 1.000 | 1.76 [1.45, 2.08], *pd* = 1.000 | −0.30 [−0.75, 0.15], *pd* = 0.911 |
| Censored mixed model (endured trials censored at ≥ 11.0 s) | −5.20 [−5.63, −4.81], *pd* = 1.000 | 2.50 [2.02, 3.01], *pd* = 1.000 | −1.46 [−2.10, −0.84], *pd* = 1.000 |

*Note.* All models: $latency \sim temperature \times instruction + (1 + temperature \times instruction \mid participant)$. Temperature and instruction are main effects averaged over the other factor, as in Table S3. Estimates in seconds [95% interval]. Primary model: lme4, REML, df and p by Satterthwaite; trials on which the participant endured the stimulus enter as 11.0 s. Censored model: those trials enter as censored (≥ 11.0 s) and effects are on the latent, uncensored latency; reporting the posterior mean, 95% credible interval and probability of direction (pd). Rows 1 and 2 fit the same model to the same data and differ only in estimation method.

**Supplementary Table S3.** Linear mixed models of effects of temperature and instruction on withdrawal latency and peak acceleration

*a. Fixed effects*

| Model | Term | Estimate | 95% CI | *df* | *t* | *F* | *p* |
|---|---|---|---|---|---|---|---|
| Withdrawal latency (s) | Intercept (grand mean) | 7.236 | [6.927, 7.545] | 90.0 | 45.86 | – | $3.33 \times 10^{-64}$ |
| | Temperature (48 − 46 °C) | −3.799 | [−4.029, −3.568] | 90.0 | −32.33 | 1045.00 | $2.57 \times 10^{-51}$ |
| | Instruction (tolerance − threshold) | 1.753 | [1.436, 2.070] | 90.0 | 10.85 | 117.67 | $5.01 \times 10^{-18}$ |
| | Temperature × instruction | −0.294 | [−0.743, 0.154] | 90.1 | −1.29 | 1.66 | .201 |
| Peak acceleration (g) | Intercept (grand mean) | 1.511 | [1.438, 1.585] | 91.1 | 40.31 | – | $6.89 \times 10^{-60}$ |
| | Temperature (48 − 46 °C) | 0.097 | [0.059, 0.135] | 90.6 | 5.05 | 25.46 | $2.31 \times 10^{-6}$ |
| | Instruction (tolerance − threshold) | −0.043 | [−0.090, 0.005] | 93.8 | −1.77 | 3.12 | .081 |
| | Temperature × instruction | 0.044 | [−0.010, 0.097] | 1361.2 | 1.60 | 2.57 | .109 |

*b. Temperature effect within each instruction*

| Model | Instruction | Temperature effect (48 − 46 °C) | 95% CI | *df* | *t* | *p* | $p_{adj}$ |
|---|---|---|---|---|---|---|---|
| Withdrawal latency (s) | Threshold | −3.651 | [−3.967, −3.336] | 90.0 | −22.98 | $2.03 \times 10^{-39}$ | $2.03 \times 10^{-39}$ |
| | Tolerance | −3.946 | [−4.282, −3.610] | 90.0 | −23.36 | $5.53 \times 10^{-40}$ | $1.11 \times 10^{-39}$ |
| Peak acceleration (g) | Threshold | 0.075 | [0.033, 0.118] | 140.8 | 3.53 | $5.57 \times 10^{-4}$ | $5.57 \times 10^{-4}$ |
| | Tolerance | 0.119 | [0.069, 0.170] | 222.3 | 4.64 | $5.98 \times 10^{-6}$ | $1.20 \times 10^{-5}$ |

*Note.* Trial-level linear mixed models, $metric \sim temperature \times instruction$, REML; df and p by Satterthwaite. Both factors coded −0.5 / +0.5, so temperature and instruction are main effects (averaged over the other factor) in the metric's units, the intercept is the grand mean, and F = $t^2$ (Type III). Random effects: Withdrawal latency (s): $(1 + Temperature * Instruction|participant)$; Peak acceleration (g): $(1 + Temperature + Instruction|participant)$. The maximal structure was fitted first and reduced along a fixed ladder only on singularity or non-convergence. Latency includes endured trials at 11.0 s (see Table S2).

**Supplementary Table S4.** Correlation between withdrawal latency and peak acceleration

| Temperature | Instruction | n | *r* [95% CI] | *p* |
|---|---|---|---|---|
| 46 °C | Threshold | 65 | .10 [−.15, .33] | .435 |
| | Tolerance | 36 | −.23 [−.52, .10] | .173 |
| 48 °C | Threshold | 90 | −.18 [−.38, .02] | .082 |
| | Tolerance | 84 | −.17 [−.37, .04] | .117 |

*Note.* Between participants: Pearson correlation of participant means. Only trials with a measured withdrawal enter, as peak acceleration is undefined for endured trials, so n is lower where many trials were endured.

**Supplementary Table S5.** Robustness of the associations with the separate pain rating

*a. 48 °C*

| Analysis | Latency, threshold | Latency, tolerance | Peak acc., threshold | Peak acc., tolerance |
|---|---|---|---|---|
| Pearson r (primary) | −.47 [−.61, −.29]<br>$p_{adj}$ = 2.08 × $10^{-5}$, n = 91 | −.50 [−.64, −.33]<br>$p_{adj}$ = 3.29 × $10^{-6}$, n = 91 | −.01 [−.22, .19]<br>$p_{adj}$ = 1.00, n = 90 | −.05 [−.26, .17]<br>$p_{adj}$ = 1.00, n = 84 |
| Partial r, controlling for withdrawal-task rating ‡ | −.49 [−.63, −.31]<br>*p* = 1.00 × $10^{-6}$, n = 91 | −.53 [−.66, −.36]<br>*p* = 8.21 × $10^{-8}$, n = 91 | .01 [−.20, .22]<br>*p* = .926, n = 90 | −.06 [−.27, .16]<br>*p* = .586, n = 84 |
| Spearman ρ | −.50 [−.65, −.32]<br>*p* = 4.03 × $10^{-7}$, n = 91 | −.54 [−.68, −.37]<br>*p* = 2.69 × $10^{-8}$, n = 91 | −.01 [−.22, .20]<br>*p* = .921, n = 90 | −.02 [−.24, .19]<br>*p* = .842, n = 84 |
| Censored mixed model, censored at 11.0 s † | −0.74 [−1.05, −0.43]<br>*pd* = 1.000, n = 91 | −1.17 [−1.64, −0.71]<br>*pd* = 1.000, n = 91 | – | – |
| Endured trials dropped | −.45 [−.60, −.27]<br>*p* = 8.70 × $10^{-6}$, n = 90 | −.47 [−.62, −.29]<br>*p* = 6.01 × $10^{-6}$, n = 84 | – | – |
| Criterion averaged across 46 and 48 °C | −.41 [−.57, −.23]<br>*p* = 4.42 × $10^{-5}$, n = 91 | −.46 [−.61, −.28]<br>*p* = 5.03 × $10^{-6}$, n = 91 | −.03 [−.24, .18]<br>*p* = .775, n = 90 | −.05 [−.26, .17]<br>*p* = .669, n = 84 |
| Criterion: arm stimuli only | −.45 [−.60, −.27]<br>*p* = 8.85 × $10^{-6}$, n = 91 | −.50 [−.64, −.33]<br>*p* = 4.31 × $10^{-7}$, n = 91 | −.05 [−.25, .16]<br>*p* = .662, n = 90 | −.10 [−.30, .12]<br>*p* = .388, n = 84 |
| Criterion: leg stimuli only | −.41 [−.57, −.22]<br>*p* = 5.83 × $10^{-5}$, n = 91 | −.41 [−.57, −.23]<br>*p* = 4.47 × $10^{-5}$, n = 91 | .02 [−.19, .23]<br>*p* = .850, n = 90 | .00 [−.21, .22]<br>*p* = .995, n = 84 |

**Supplementary Table S5** (continued)

*b. 46 °C*

| Analysis | Latency, threshold | Latency, tolerance | Peak acc., threshold | Peak acc., tolerance |
|---|---|---|---|---|
| Pearson r (primary) | −.36 [−.53, −.17]<br>$p_{adj}$ = .002, n = 91 | −.32 [−.49, −.12]<br>$p_{adj}$ = .010, n = 91 | −.07 [−.30, .18]<br>$p_{adj}$ = 1.00, n = 65 | .08 [−.25, .40]<br>$p_{adj}$ = 1.00, n = 36 |
| Partial r, controlling for withdrawal-task rating ‡ | −.25 [−.44, −.05]<br>*p* = .015, n = 91 | −.24 [−.43, −.04]<br>*p* = .022, n = 91 | −.09 [−.33, .16]<br>*p* = .458, n = 65 | .06 [−.28, .38]<br>*p* = .741, n = 36 |
| Spearman ρ | −.39 [−.55, −.19]<br>*p* = 1.51 × $10^{-4}$, n = 91 | −.33 [−.51, −.13]<br>*p* = .001, n = 91 | −.27 [−.49, −.03]<br>*p* = .027, n = 65 | −.12 [−.44, .22]<br>*p* = .475, n = 36 |
| Censored mixed model, censored at 11.0 s † | −1.12 [−1.70, −0.55]<br>*pd* = 1.000, n = 91 | −1.03 [−1.67, −0.40]<br>*pd* = 0.999, n = 91 | – | – |
| Endured trials dropped | −.30 [−.51, −.06]<br>*p* = .015, n = 65 | −.22 [−.51, .12]<br>*p* = .197, n = 36 | – | – |
| Criterion averaged across 46 and 48 °C | −.41 [−.57, −.22]<br>*p* = 6.17 × $10^{-5}$, n = 91 | −.36 [−.53, −.17]<br>*p* = 4.28 × $10^{-4}$, n = 91 | −.06 [−.30, .18]<br>*p* = .621, n = 65 | .09 [−.24, .41]<br>*p* = .582, n = 36 |
| Criterion: arm stimuli only | −.35 [−.52, −.16]<br>*p* = 5.78 × $10^{-4}$, n = 91 | −.36 [−.52, −.16]<br>*p* = 5.44 × $10^{-4}$, n = 91 | −.08 [−.32, .16]<br>*p* = .503, n = 65 | .09 [−.25, .40]<br>*p* = .616, n = 36 |
| Criterion: leg stimuli only | −.32 [−.49, −.12]<br>*p* = .002, n = 91 | −.24 [−.42, −.03]<br>*p* = .024, n = 91 | −.04 [−.28, .20]<br>*p* = .733, n = 65 | .07 [−.27, .39]<br>*p* = .700, n = 36 |

*Note.* Cells: estimate [95% CI]; below it the p value (unadjusted: these are sensitivity checks of the primary tests, whose Holm-adjusted values are in the main text), or pd for the Bayesian censored model row, and n participants. Pearson: primary analysis, participant means with endured trials at 11.0 s. ‡ Partial correlation of the same participant means, controlling for the participant's mean pain rating given after each trial of the withdrawal task (CI by Fisher z). Spearman CIs use the Bonett–Wright standard error. † Censored regression on trials, with endured trials right-censored at ≥ 11.0 s. Latency was regressed on the separate thermal rating task, with a separate rating slope and residual SD for each instruction × temperature condition and uncorrelated by-participant random intercepts per condition. The model was fitted in brms with the same settings as the censored model in Table S2 (posterior mean [95% credible interval]; pd = probability of direction). The estimate is the change in latent withdrawal latency (s) for a 10-point higher rating in the separate experiment (negative = earlier withdrawal). Endured trials dropped: participant means over measured trials only (at least 3). Criterion averaged: one separate-experiment rating per participant over both temperatures. Arm / leg: temperature-matched rating over stimuli to that site only (at least 6). Censoring does not apply to peak acceleration.
